## Supporting Text for "Herpes simplex virus 2 (HSV-2) evolves faster in cell culture than HSV-1 by generating greater genetic diversity"

**S1 Text. Supporting Material and Methods for HSV replication kinetics and infection models.**

**Virus growth curves**

Vero cell monolayers with 1 x 10^5^ cells/well in 12-well plates were infected with the indicated virus for 1 h, at high multiplicity of infection (MOI), i.e. 5 PFU/cell, or low MOI (0.01 PFU/cell) for the one-step and multi-step growth curves, respectively. Cells were then washed and fresh 2% FBS DMEMs was added. At indicated times post-infection, the medium was harvested and centrifuged at 500 x g for 5 min to pellet detached cells. These cells were combined with infected cells that had been scraped from the well into 0.5 ml of fresh medium. Samples were frozen/thawed three times and titrated on Vero cells in triplicates.

**Ethical statement**

Animal care and sacrifice were performed in strict accordance with the European Health Law of the Federation of Laboratory Animal Science Association and the Spanish Animal Welfare Law. All animal experiments were performed with special efforts to minimize animal suffering, in compliance with national and international regulations and were approved by the Ethical Review Board of Consejo Superior de Investigaciones Científicas and Comunidad de Madrid under reference PROEX 025/16. The total number of used animals was reported at the end of each year to the animal welfare deputy of the Animal Facility and Comunidad de Madrid.

**Infection and pathogenesis monitoring of mice**

BALB/cByJ female 5-8 weeks old mice were purchased from Charles Rivers. The infections of groups of five animals per condition were performed with viral stocks diluted in PBS containing 0.1% bovine serum albumin. Mice were anaesthetized with isoflurane and then infected by intranasal (i.n.) inoculation with 10 µl of virus inoculum containing the indicated infectious doses. On the other hand, mice were pretreated subcutaneously with 2.5 mg of medroxyprogesterone acetate (Pfizer) diluted in 100 µl of PBS. Five days later, mice were anaesthetized with isoflurane and then infected by intravaginal (i.v.) inoculation with 10 µl of viral inoculum containing the indicated infectious doses. Viral doses were confirmed by titrating again the virus dilutions used for mouse infections, by plaque assay on Vero cells. Mice were housed in ventilated racks and monitored daily for survival, weight and signs of illness. Disease for i.n. infections was scored as follows: (1) ruffled fur, (2) reduced mobility / moderate blindness / stooped, (3) breathing difficulties / severe blindness / generalized morbidity, and (4, humane endpoint) severe breathing difficulties / convulsions / forelimb paralysis (one leg). Infections through the i.v. route were scored as (1) ruffled fur, (2) reduced mobility / genital swelling / stooped, (3) generalized morbidity / moderate abdominal swelling, and (4, humane endpoint) anal occlusion or evident constipation / severe abdominal swelling / hind limb paralysis (one leg).

**Statistical analysis**

Data analysis was performed using GraphPad Prism 8 (v8.4.3) software. Percentage of initial weight data were analyzed using multiple *t*-tests with Sidak-Bonferroni correction (*p* < 0.05). Analyses were performed up to times post-infection at which survival rates in the corresponding groups were above 50%.
