## Supporting Figures for "Herpes simplex virus 2 (HSV-2) evolves faster in cell culture than HSV-1 by generating greater genetic diversity"

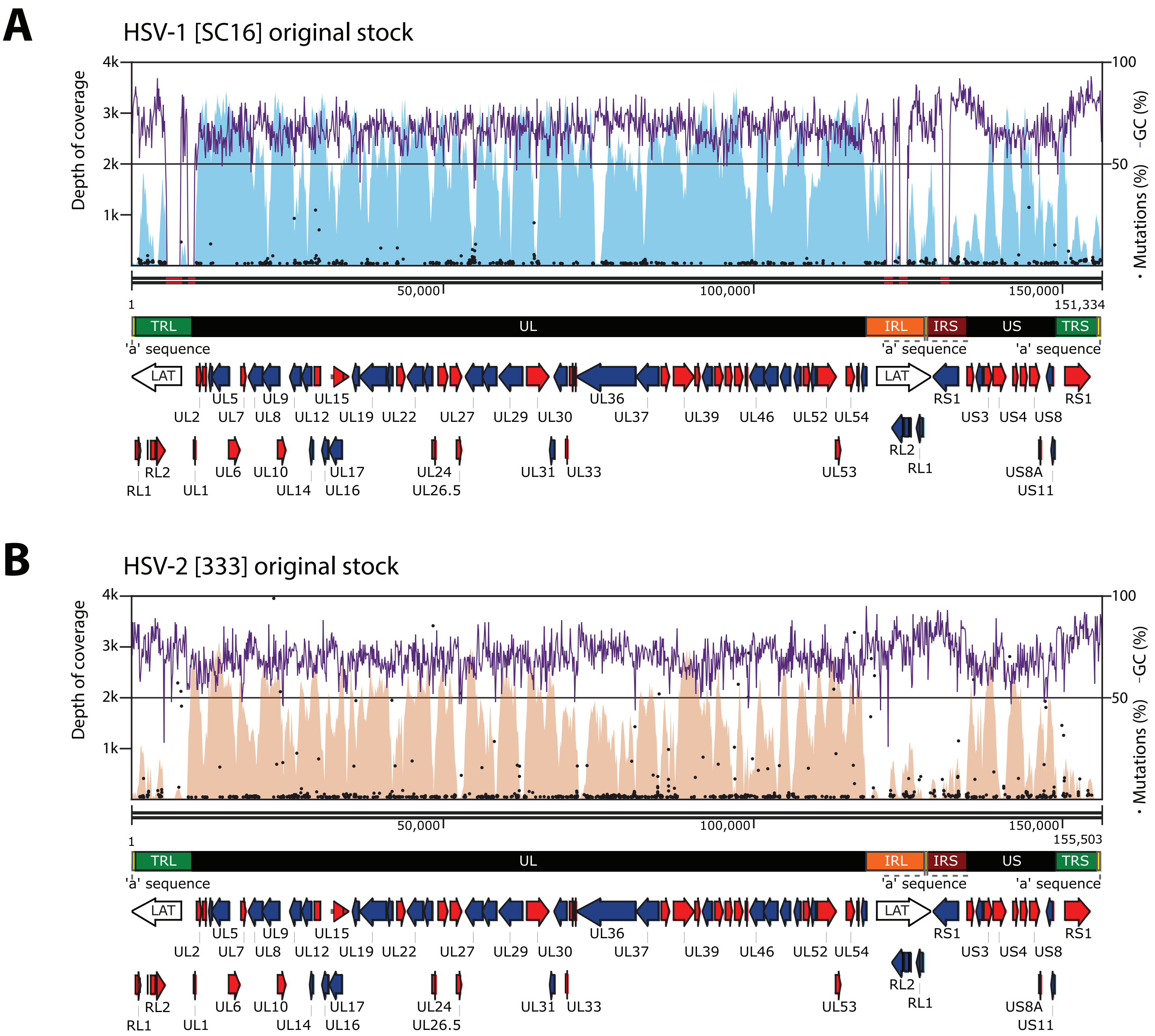


**S1 Fig. Schematic of the HSV-1 strain SC16 (A) and HSV-2 strain 333 (B) sequenced genomes from original stocks.** Each CDS is presented in forward (red) or reverse (blue) orientation. Detected MVs are mapped as black (not de novo) or red (de novo) dots across the genome, according to their location (*x*-axis) and frequency (*y*-axis). GC% plots (purple lines) and coverage plots from data alignments (blue/orange profiles) have also been mapped across each genome.

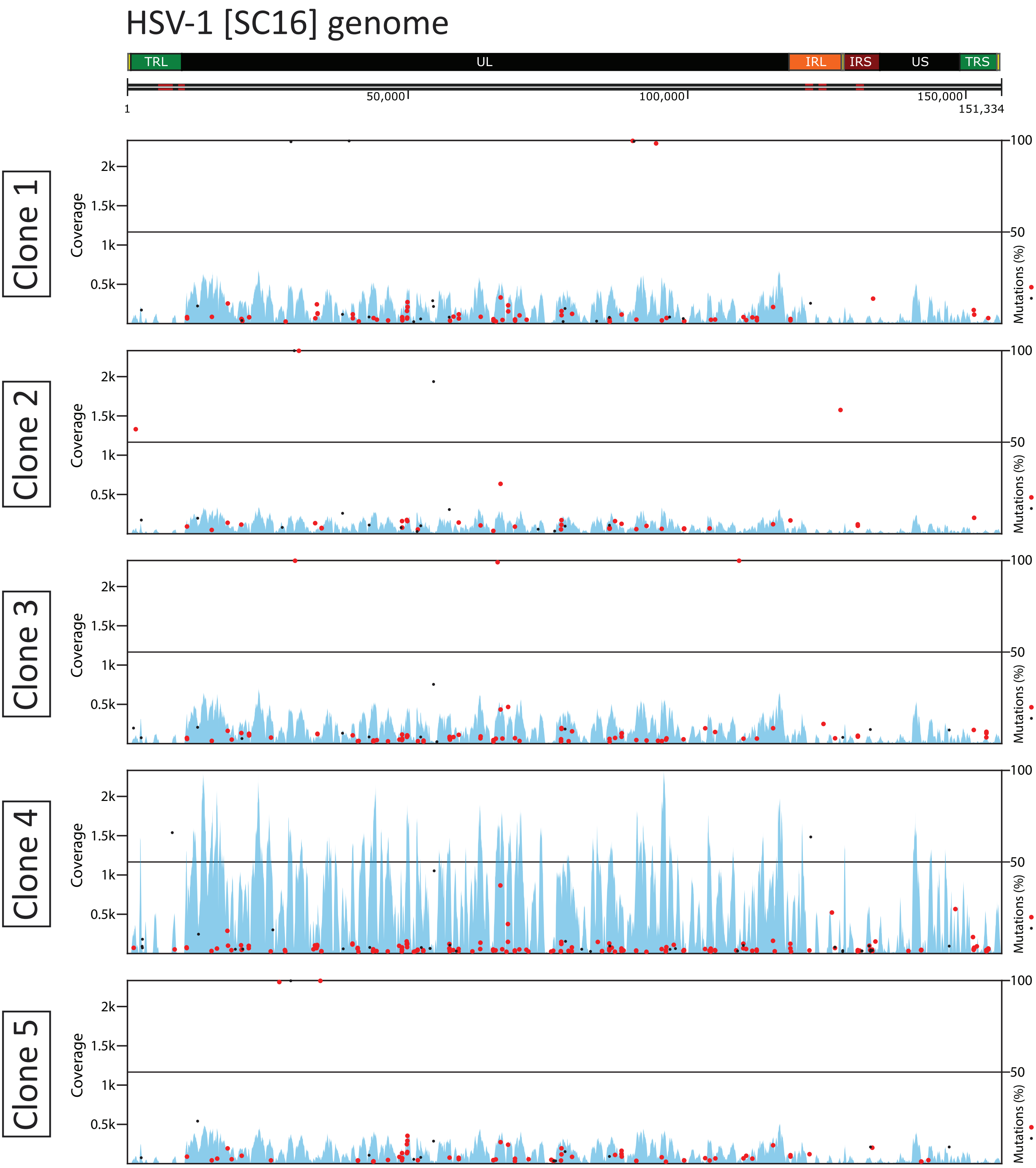


**S2 Fig. Variant analysis of HSV-1 plaque-purified clones.** Coverage plots from data alignments are represented in blue, for each individual case. Detected MVs are mapped as black (not de novo) or red (de novo) dots across the genome, according to their location (*x*-axis) and frequency (*y*-axis). MVs were considered as de novo when these were not previously found in the original stock (see Materials and Methods for details).

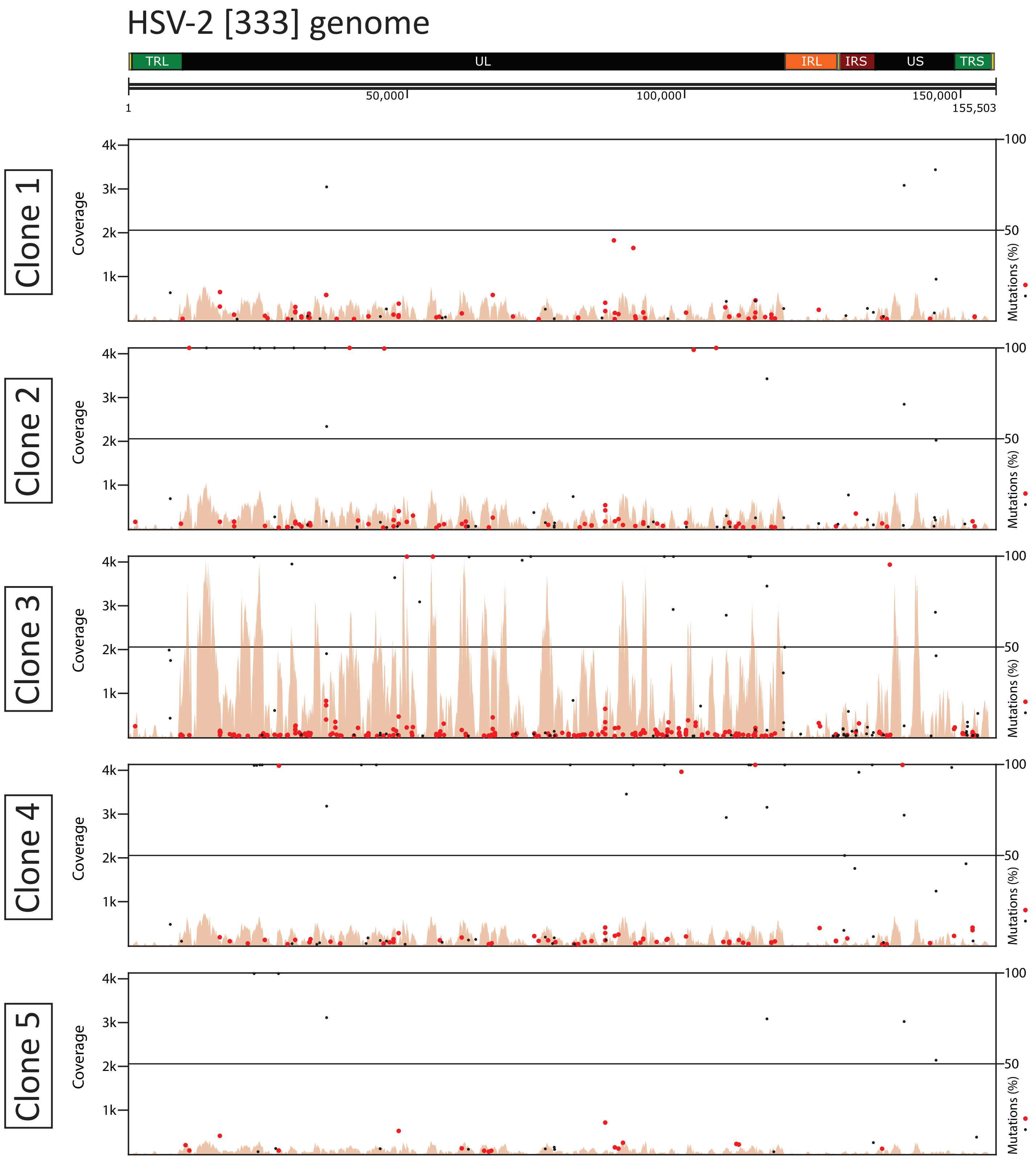


**S3 Fig. Variant analysis of HSV-2 plaque-purified clones.** Coverage plots from data alignments are represented in orange, for each individual case. Detected MVs are mapped as black (not de novo) or red (de novo) dots across the genome, according to their location (*x*-axis) and frequency (*y*-axis). MVs were considered as de novo when these were not previously found in the original stock.

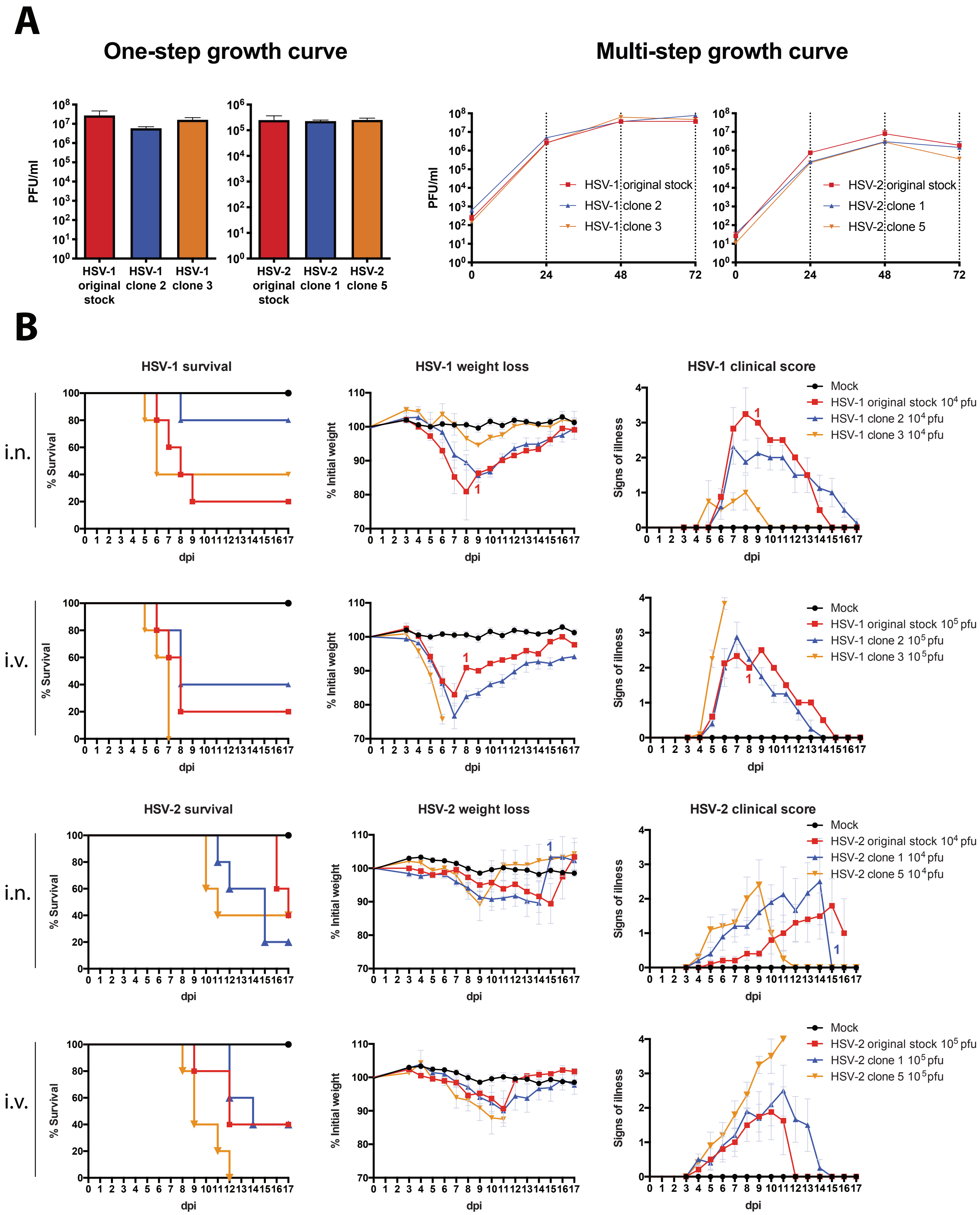


**S4 Fig. Replication kinetics and pathogenesis of HSV-1 and HSV-2 plaque-isolated clones compared to their corresponding original stocks.** (A) Vero cells were infected with the indicated viruses at high MOI (5 PFU/cell) for one-step growth curves, and at low MOI (0.01 PFU/cell) for multi-step growth curves. Virus titers from fractions containing cell-associated virus were determined by plaque assay at 24 hpi in the one-step curves, and at the indicated times in the multi-step curves. (B) Female BALB/c mice (n=5) were infected with the indicated virus and dose, by intranasal (i.n.) or intravaginal (i.v.) inoculations. Mice were monitored daily for survival, body weight, and signs of illness. Weight data are expressed as the mean +/- SEM of the five animal weights compared to their original weight on the day of inoculation. Signs of illness, as a score ranged from 1 to 4, is also expressed as the mean +/- SEM of the five animals. A colored “1” indicates thereafter only one animal remained in that group. Statistical analysis was performed for bodyweight data, using multiple *t*-tests with Sidak-Bonferroni correction ($p < 0.05$).

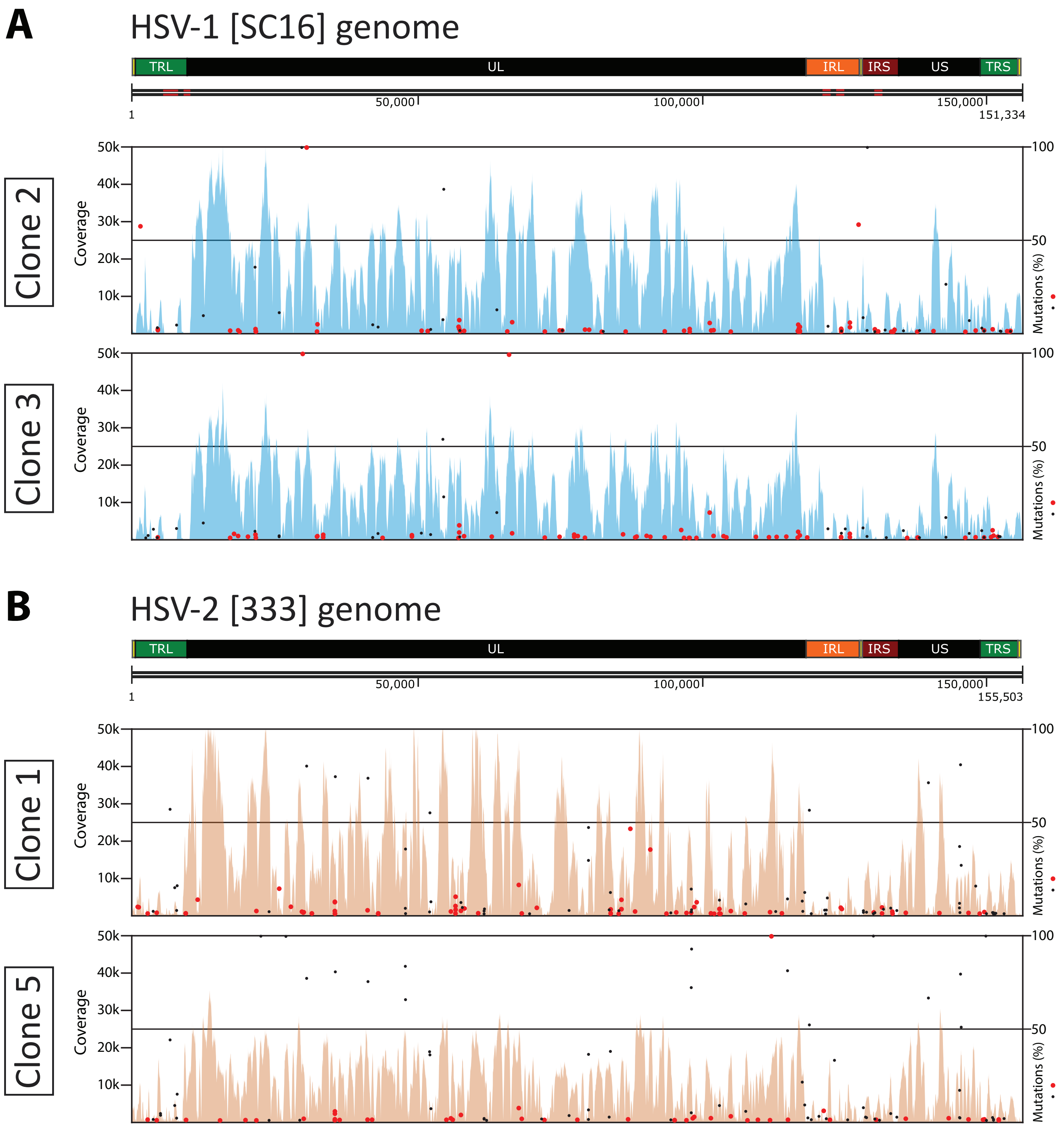


**S5 Fig. Variant analysis of HSV-1 plaque-purified clones 2 and 3 (A) and HSV-2 clones 1 and 5 (B) from high-depth sequencing data.** Coverage plots from alignments are represented in blue or orange, for each case. Detected MVs are mapped as black (not de novo) or red (de novo) dots across the genome, according to their location (*x*-axis) and frequency (*y*-axis). MVs were considered as de novo when these were not previously found in the corresponding original stock.

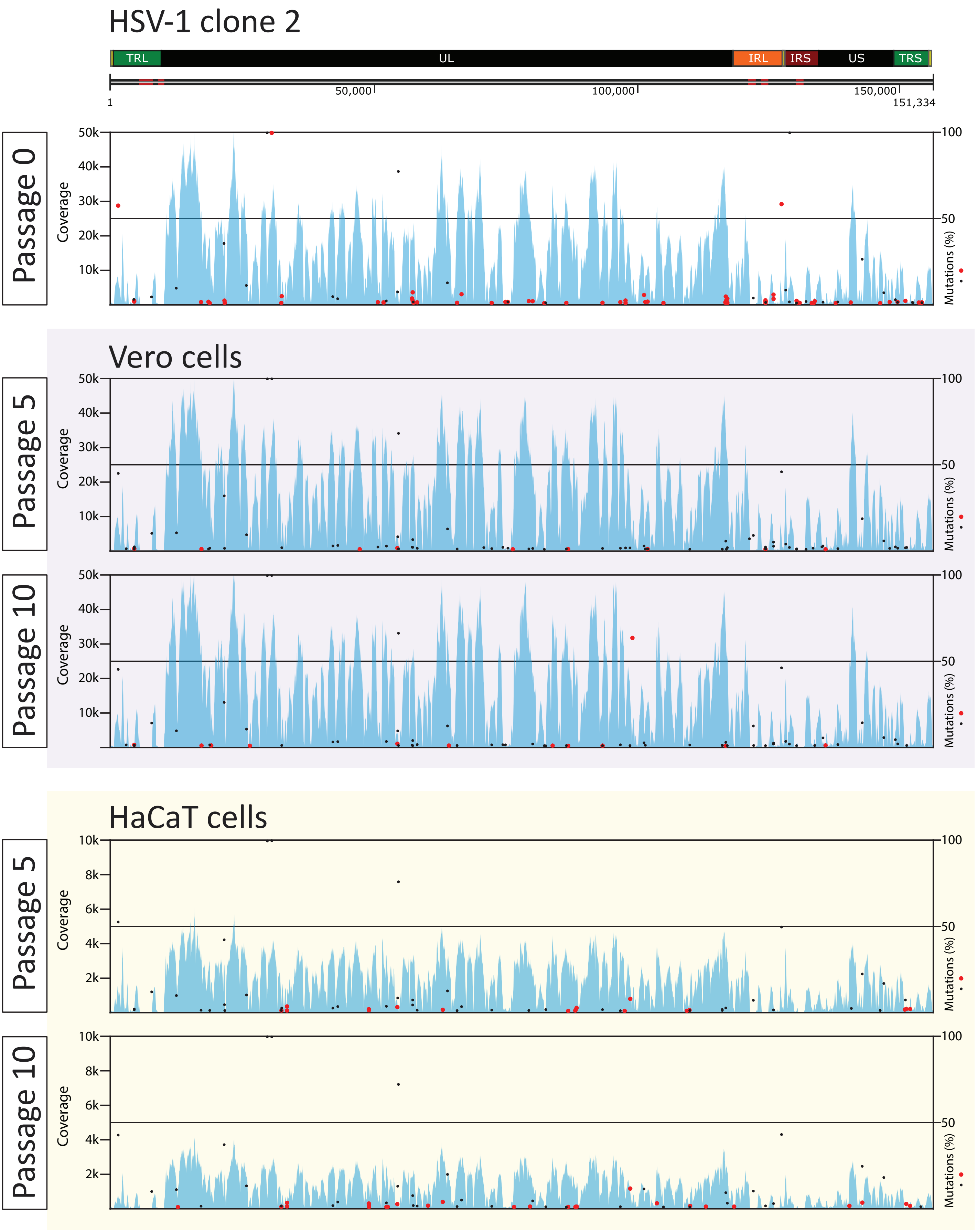


**S6 Fig. Variant analysis of HSV-1 plaque-purified clone 2, after 5 and 10 passages in Vero and HaCaT cells.** Coverage plots from high-depth sequencing data alignments are represented in blue. Detected MVs are mapped as black (not *de novo*) or red (*de novo*) dots across the genome, according to their location (*x*-axis) and frequency (*y*-axis). Mutations from passage 0 were considered as de novo when these were not previously found in the original stock, whereas those from passage 5 and 10, regarding passage 0.

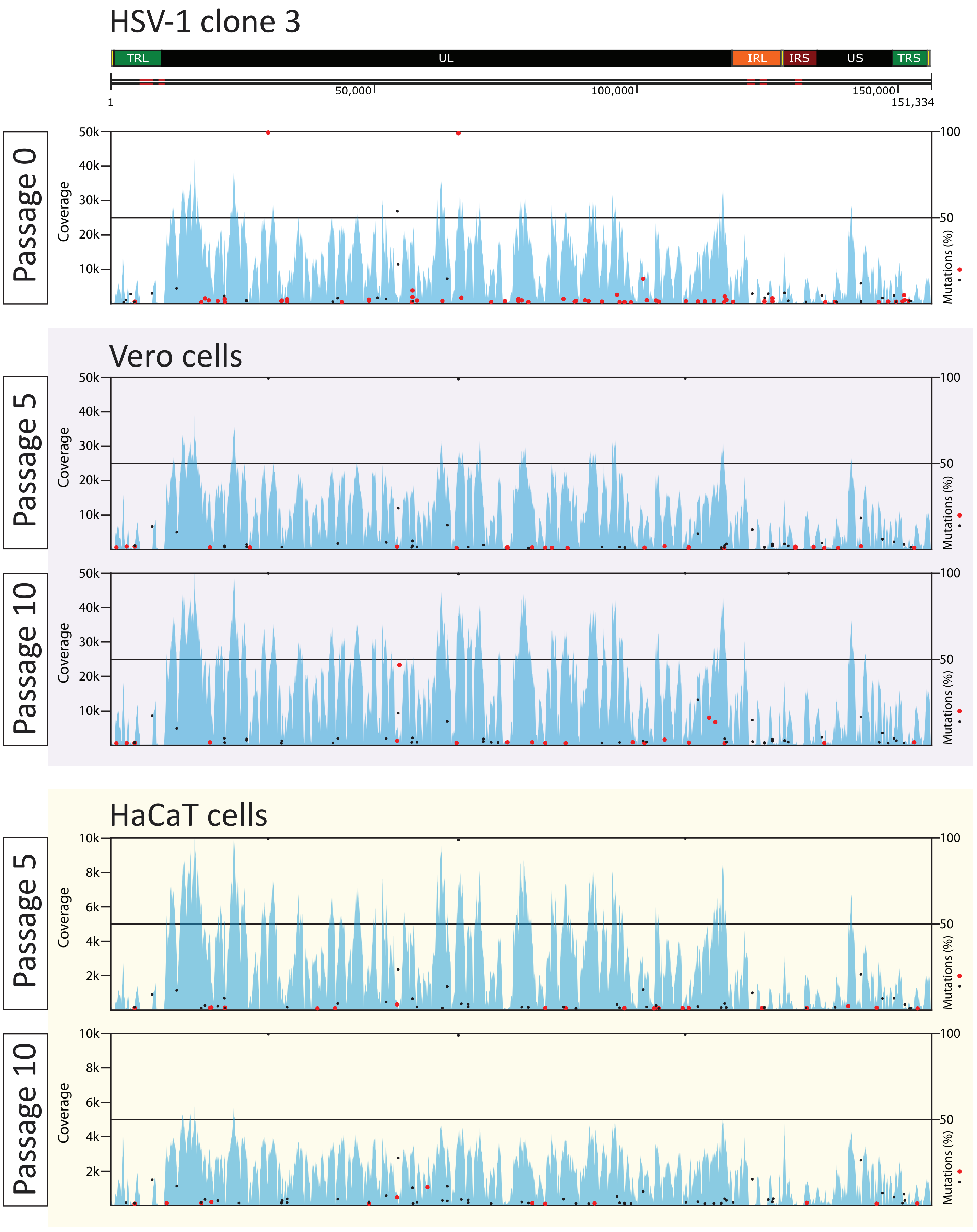


**S7 Fig. Variant analysis of HSV-1 plaque-purified clone 3, after 5 and 10 passages in Vero and HaCaT cells.** Coverage plots from high-depth sequencing data alignments are represented in blue. Detected MVs are mapped as black (not *de novo*) or red (*de novo*) dots across the genome, according to their location (*x*-axis) and frequency (*y*-axis). Mutations from passage 0 were considered as de novo when these were not previously found in the original stock, whereas those from passage 5 and 10, regarding passage 0.

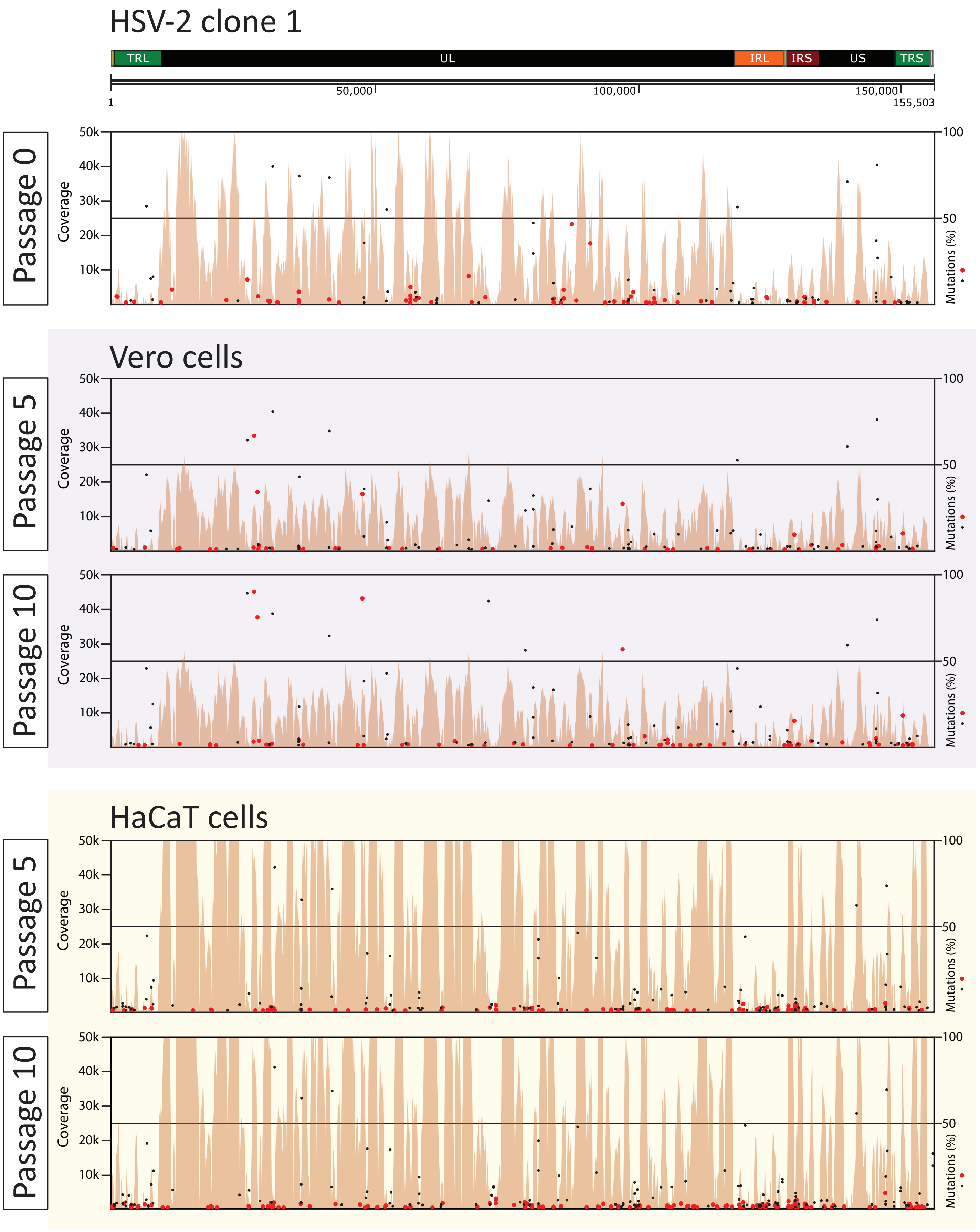


**S8 Fig. Variant analysis of HSV-2 plaque-purified clone 1, after 5 and 10 passages in Vero and HaCaT cells.** Coverage plots from high-depth sequencing data alignments are represented in orange. Detected MVs are mapped as black (not *de novo*) or red (*de novo*) dots across the genome, according to their location (*x*-axis) and frequency (*y*-axis). Mutations from passage 0 were considered as de novo when these were not previously found in the original stock, whereas those from passage 5 and 10, regarding passage 0.

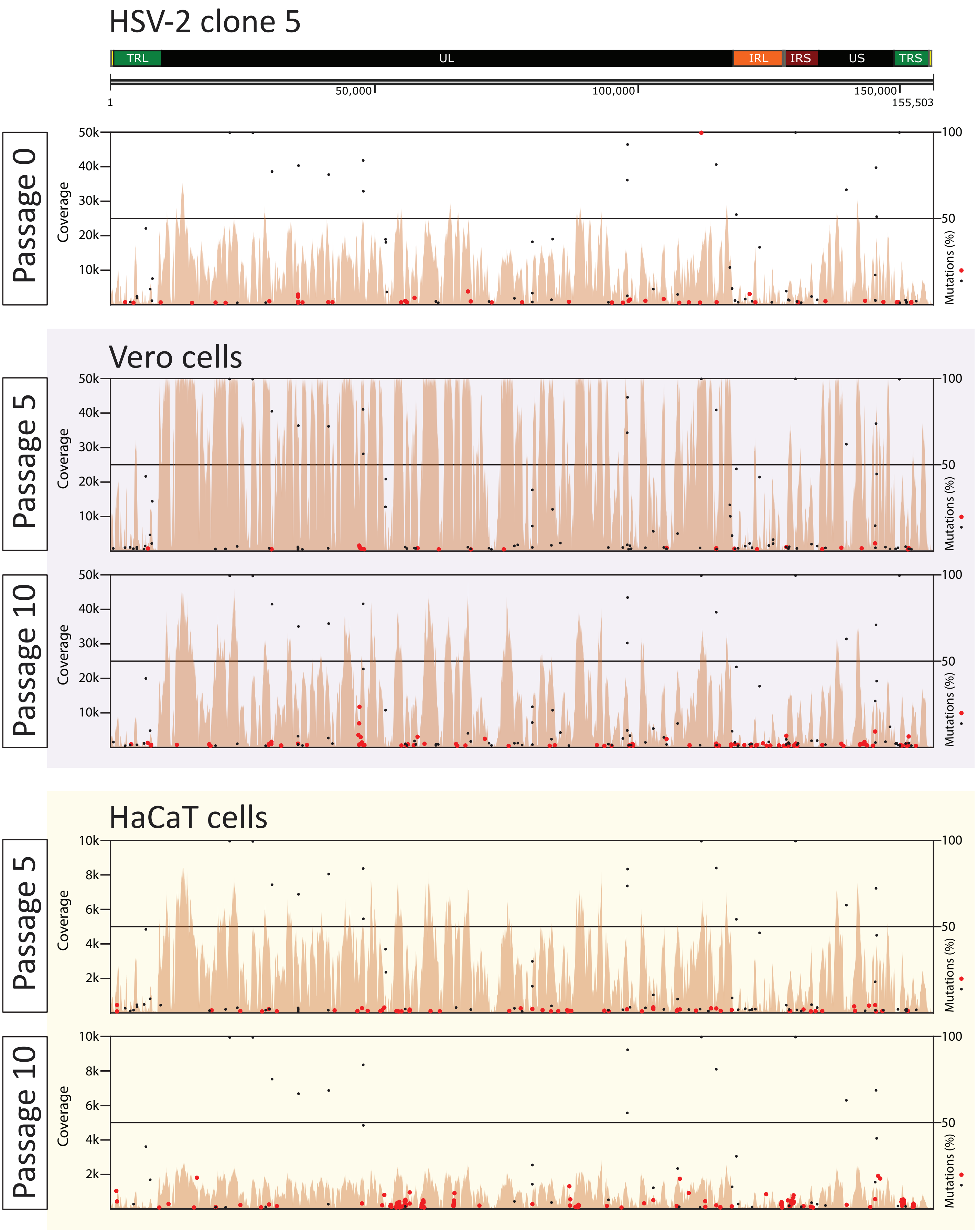


**S9 Fig. Variant analysis of HSV-2 plaque-purified clone 5, after 5 and 10 passages in Vero and HaCaT cells.** Coverage plots from high-depth sequencing data alignments are represented in orange. Detected MVs are mapped as black (not *de novo*) or red (*de novo*) dots across the genome, according to their location (*x*-axis) and frequency (*y*-axis). Mutations from passage 0 were considered as de novo when these were not previously found in the original stock, whereas those from passage 5 and 10, regarding passage 0.
