## Supplementary figures and images for "Herpes simplex virus 2 (HSV-2) evolves faster in cell culture than HSV-1 by generating greater genetic diversity"

### Supporting Animation 1

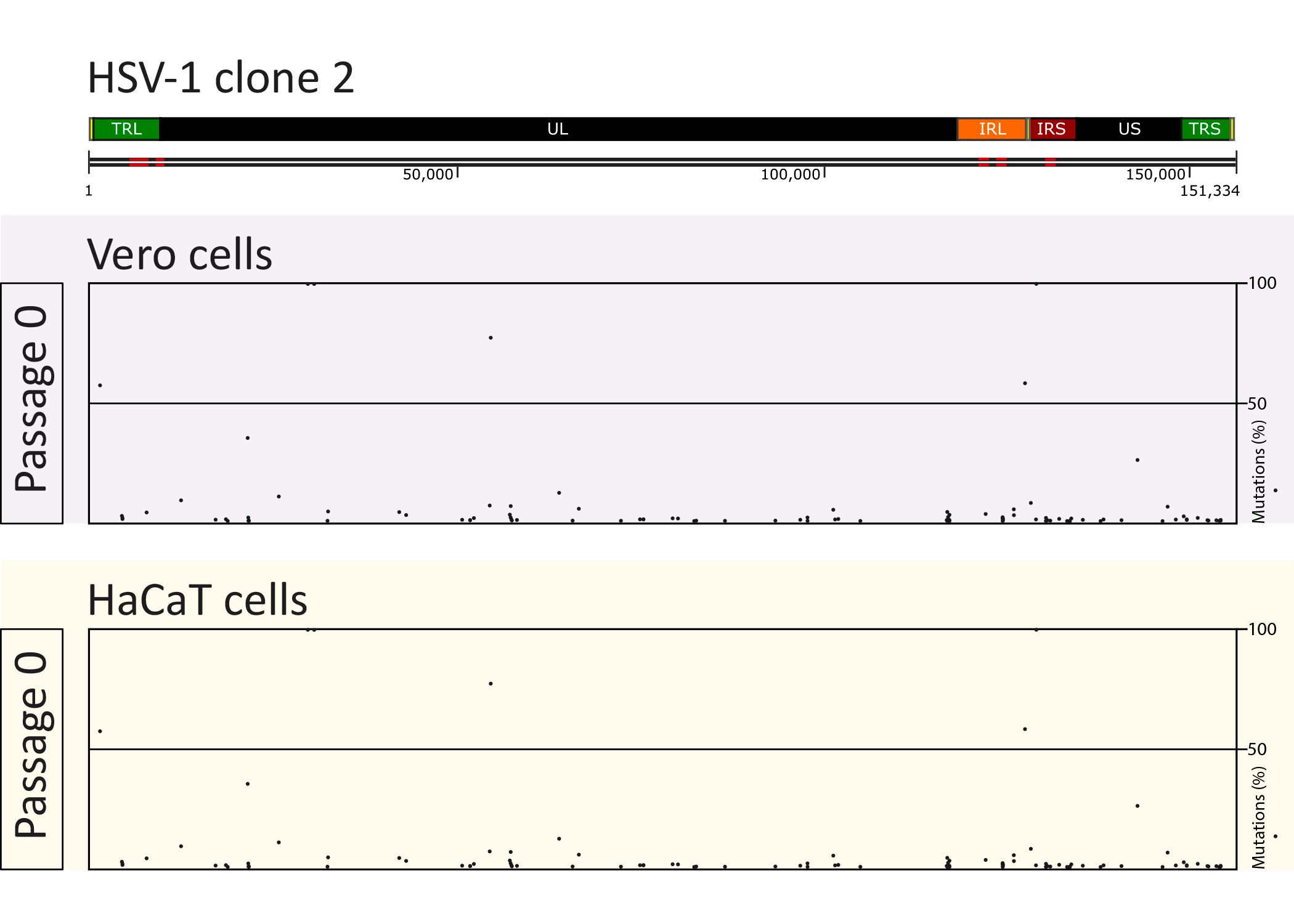

### Supporting Animation 2

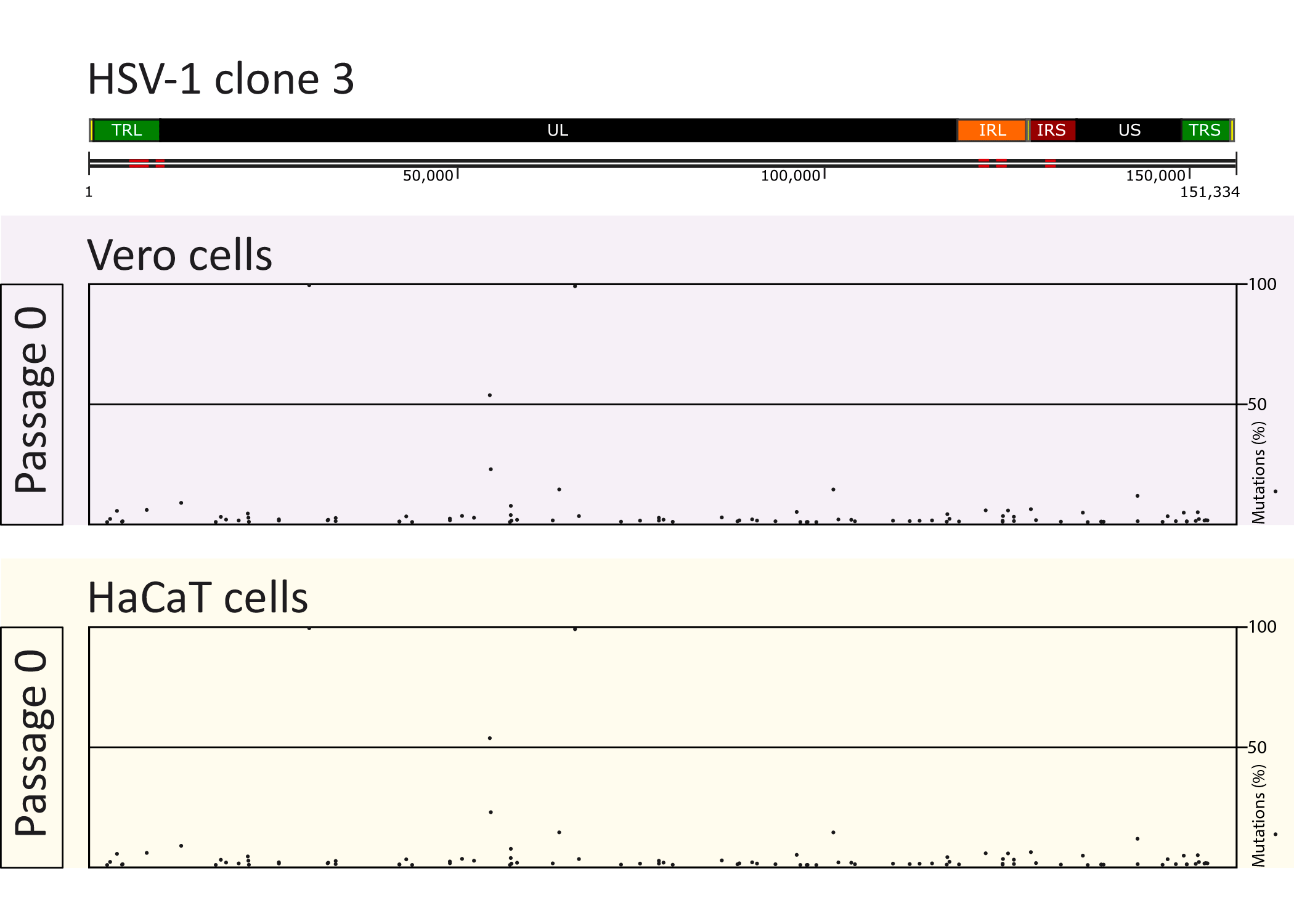

### Supporting Animation 3

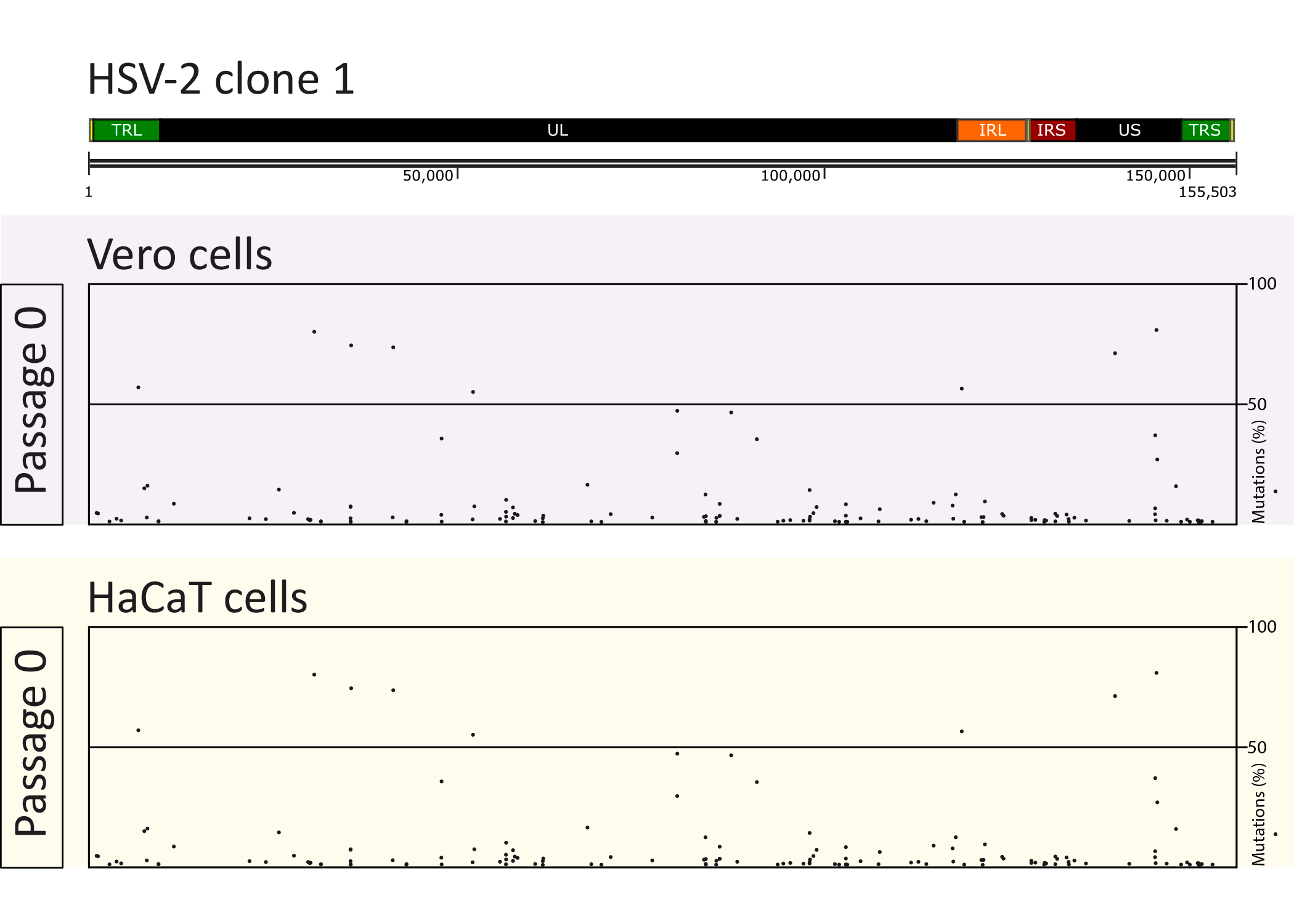

### Supporting Animation 4

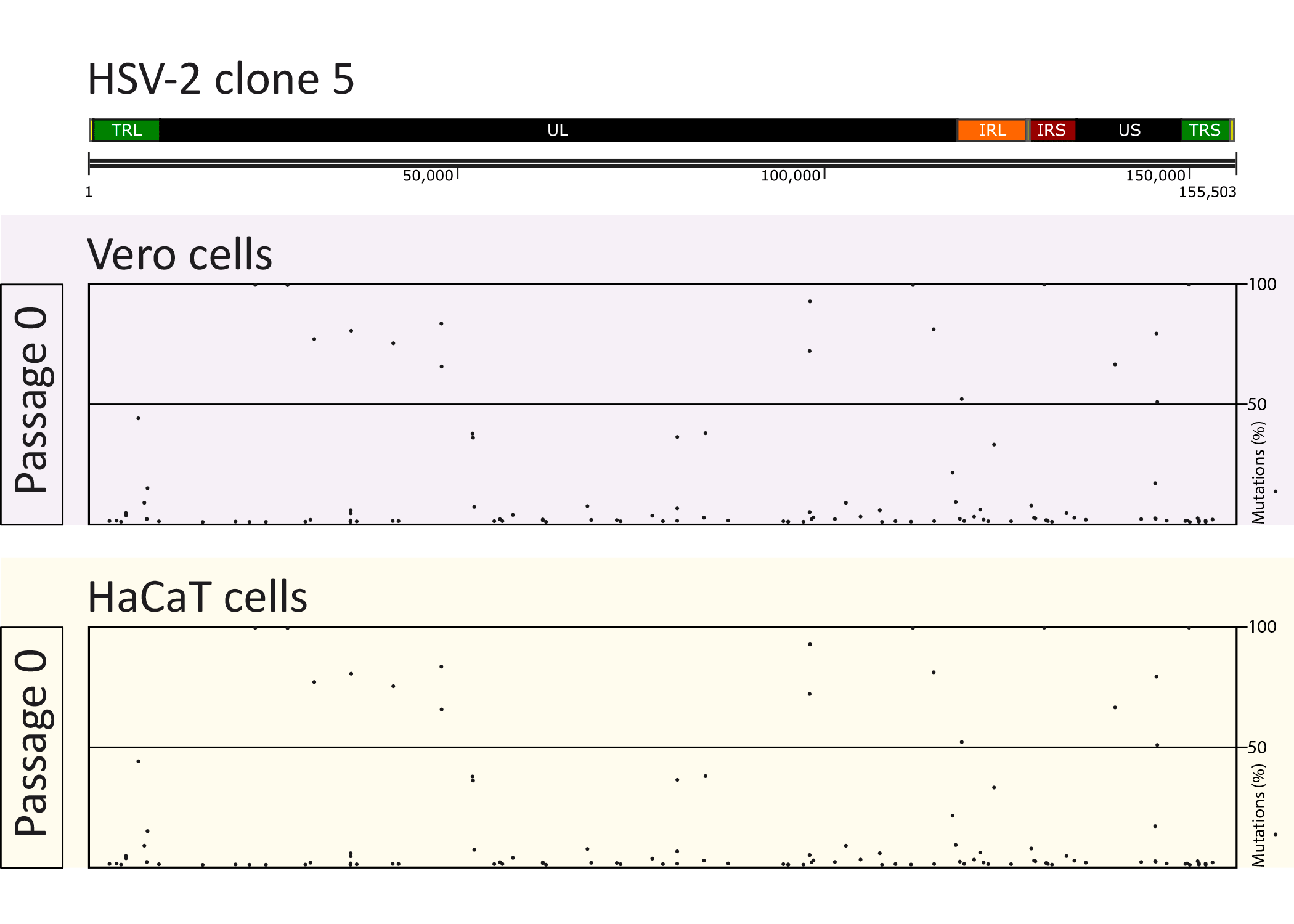
